# The origin and underlying driving forces of the SARS-CoV-2 outbreak

**DOI:** 10.1101/2020.04.12.038554

**Authors:** Shu-Miaw Chaw, Jui-Hung Tai, Shi-Lun Chen, Chia-Hung Hsieh, Sui-Yuan Chang, Shiou-Hwei Yeh, Wei-Shiung Yang, Pei-Jer Chen, Hurng-Yi Wang

**Author notes:** Corresponding Author: (HYW). These authors contributed equally to this work (Shu-Miaw Chaw, Hurng-Yi Wang).

## Abstract

The spread of SARS-CoV-2 since December 2019 has become a pandemic and impacted many aspects of human society. Here, we analyzed genetic variation of SARS-CoV-2 and its related coronavirus and found the evidence of intergenomic recombination. After correction for mutational bias, analysis of 137 SARS-CoV-2 genomes as of 2/23/2020 revealed the excess of low frequency mutations on both synonymous and nonsynonymous sites which is consistent with recent origin of the virus. In contrast to adaptive evolution previously reported for SARS-CoV in its brief epidemic in 2003, our analysis of SARS-CoV-2 genomes shows signs of relaxation of selection. The sequence similarity of the spike receptor binding domain between SARS-CoV-2 and a sequence from pangolin is probably due to an ancient intergenomic introgression. Therefore, SARS-CoV-2 might have cryptically circulated within humans for years before being recently noticed. Data from the early outbreak and hospital archives are needed to trace its evolutionary path and reveal critical steps required for effective spreading. Two mutations, 84S in orf8 protein and 251V in orf3 protein, occurred coincidentally with human intervention. The 84S first appeared on 1/5/2020 and reached a plateau around 1/23/2020, the lockdown of Wuhan. 251V emerged on 1/21/2020 and rapidly increased its frequency. Thus, the roles of these mutations on infectivity need to be elucidated. Genetic diversity of SARS-CoV-2 collected from China was two time higher than those derived from the rest of the world. In addition, in network analysis, haplotypes collected from Wuhan city were at interior and have more mutational connections, both of which are consistent with the observation that the outbreak of cov-19 was originated from China.

**SUMMARY:** In contrast to adaptive evolution previously reported for SARS-CoV in its brief epidemic, our analysis of SARS-CoV-2 genomes shows signs of relaxation of selection. The sequence similarity of the spike receptor binding domain between SARS-CoV-2 and a sequence from pangolin is probably due to an ancient intergenomic introgression. Therefore, SARS-CoV-2 might have cryptically circulated within humans for years before being recently noticed. Data from the early outbreak and hospital archives are needed to trace its evolutionary path and reveal critical steps required for effective spreading. Two mutations, 84S in orf8 protein and 251V in orf3 protein, occurred coincidentally with human intervention. The 84S first appeared on 1/5/2020 and reached a plateau around 1/23/2020, the lockdown of Wuhan. 251V emerged on 1/21/2020 and rapidly increased its frequency. Thus, the roles of these mutations on infectivity need to be elucidated.

## INTRODUCTION

A newly emerging coronavirus was detected in patients during an outbreak of respiratory illnesses starting in mid-December of 2019 in Wuhan, the capital of Hubei Province, China [1, 2, 3]. Due to the similarity of its symptoms to those induced by the severe acute respiratory syndrome (SARS) and genome organization similarity, the causal virus was named SARS-CoV-2 by the International Committee on Taxonomy of Viruses [4]. As of 3/16/2020, 167,515 cases of SARS-CoV-2 infection have been confirmed in 114 countries, causing 6,606 fatalities. As a result, WHO declared the first pandemic caused by a coronavirus on 3/11/2020 (https://www.who.int/emergencies/diseases/novel-coronavirus-2019/situation-reports). As the virus continues to spread, numerous strains have been isolated and sequenced. On 3/18/2020, more than 500 complete or nearly complete genomes have been sequenced and made publicly available.

SARS-CoV-2 is the seventh coronavirus found to infect humans. Among the other six, SARS-CoV and MERS-CoV can cause severe respiratory illness, whereas 229E, HKU1, NL63, and OC43 produce mild symptoms [5]. Current evidence strongly suggests that all human associated coronaviruses originated from other animals, such as bats and rodents [5, 6]. While SARS-CoV-2 shares similar genomic structure with other coronaviruses [7-10], its sequence differs substantially from some of the betacoronaviruses that infect humans, such as SARS-CoV (approximately 76% identity), MERS-CoV (43% identity), and HKU-1 (33% identity), but exhibits 96% similarity to a coronavirus collected in Yunnan Province, China from a bat, *Rhinolophus affinis*. Therefore, SARS-CoV-2 most likely originated from bats [2, 11].

Several issues concerning the origin, time of virus introduction to humans, evolutionary patterns, and the underlying driving force of the SARS-CoV-2 outbreak remain to be clarified [12, 13]. Here, we analyzed genetic variation of SARS-CoV-2 and its related coronaviruses. We discuss how mutational bias influences genetic diversity of the virus and attempt to infer forces that shape SARS-CoV-2 evolution.

## RESULTS

### Molecular evolution of SARS-COV-2 and related coronaviruses

The resulting phylogeny reveals that RaTG13 is the closest relative of SARS-COV-2, followed by pangolin_2019 and pangolin_2017, then CoVZC45 and CoVZXC21, and other SARS-related sequences as outgroups (S1 Fig). According to general time reversible model, transition occurred more frequent than transversion with C-T and A-G changes account for 45% and 28%, respectively, of all six types of nucleotide changes. We next estimated the strength of selection for each coding region using the dN and dS. While purifying selection tends to remove amino acid-altering mutations, thus reducing dN and dN/dS, positive selection has the opposite effect, increasing dN and dN/dS [14]. Between SARS-CoV-2 and RaTG13, *orf8* gene exhibits the highest dN (0.032) followed by *spike* (0.013) and *orf7* (0.011), all above the genome average of 0.007 (Table 1). dS varies greatly among CDSs with the highest of 0.313 in *spike* and the lowest of 0.018 in *envelope* (genome average 0.168). Finally, dN/dS is the highest in *orf8* (0.105) followed by *orf7* (0.061) and *orf3* (0.060), with the genome average of 0.042. Since *spike* shows both high dS and dN, its protein evolution rate (dN/dS) is only 0.040. Thus, while the coronavirus evolved very rapidly, it has actually been under tremendous selective constraint [13].

**Table 1.**
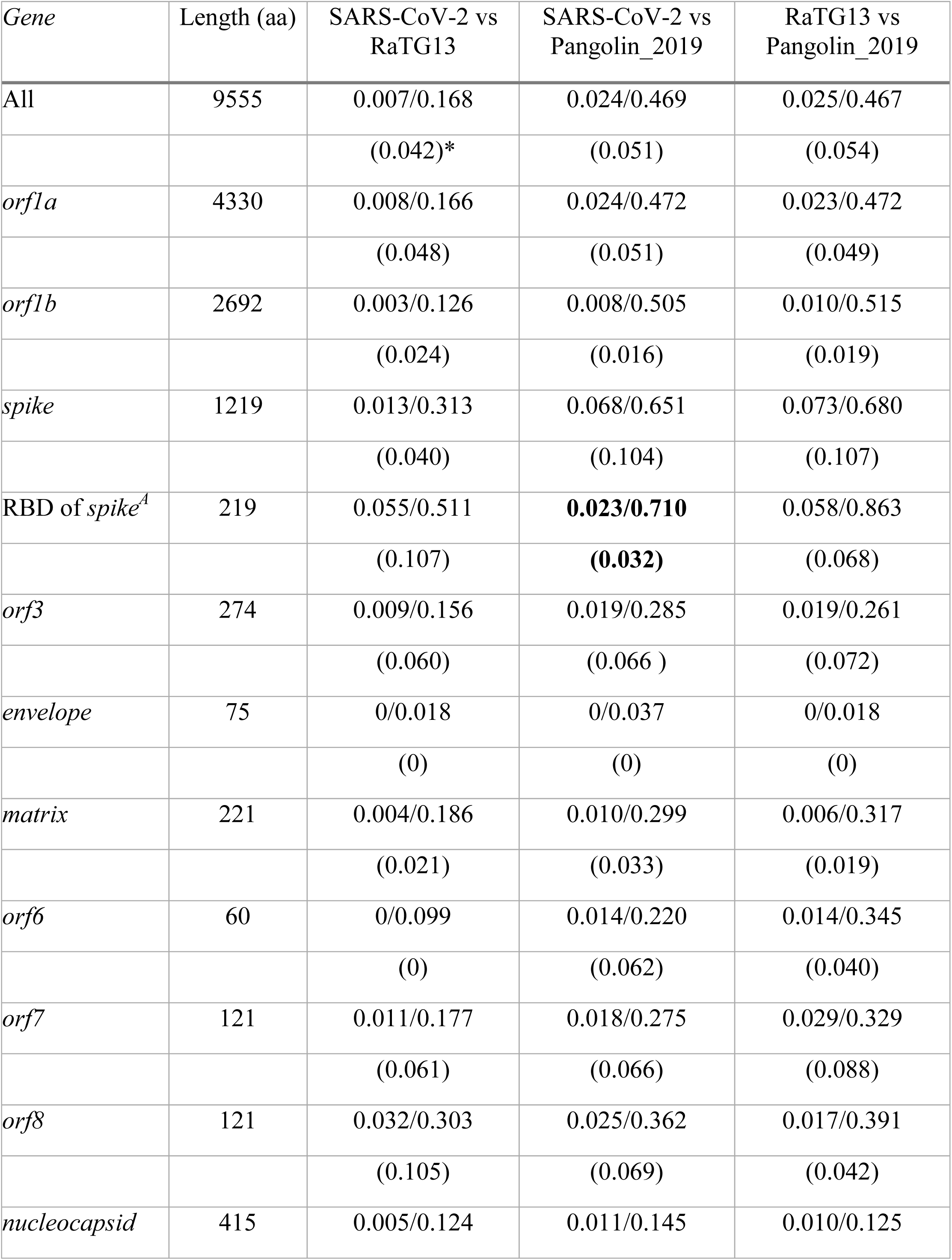
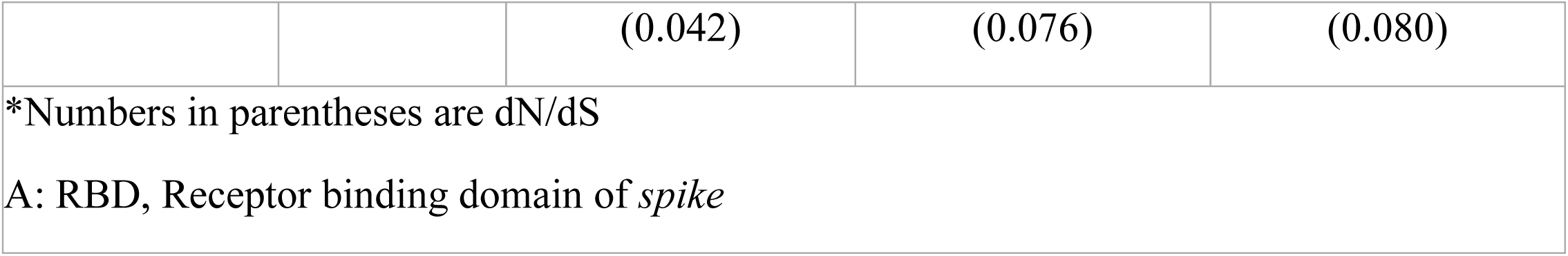
Pairwise comparison of nonsynonymous (dN; above slash) and synonymous (dS; below slash) divergence between SARS-CoV-2, RaTG13, and Pangolin_2019 of different coding regions.

Spike protein similarity between SARS-CoV-2 and pangolin_2019 led to the idea that the receptor binding domain (RBD) within the SARS-CoV-2 spike protein originated from pangolin_2019 via recombination [15-18]. If that were the case, we would expect the divergence at synonymous sites (dS) to also be reduced in the RBD region. However, while dN in the RBD region is 0.023, approximately one third of the estimate for the rest of the *spike* gene (0.068), dS in the RBD (0.710) is actually slightly higher than in the rest of the *spike* sequence (0.651). This argues against the recombination scenario. We noticed that the dS of the whole *spike* and the RBD, are 2- and 3-fold, respectively, higher than the genome average. Since synonymous sites are typically less influenced by selection, the increased divergence in dS may reflect an underlying elevated mutation rate.

### Genetic variation of SARS-CoV-2

We downloaded 137 SARS-CoV-2 genomes available from GISAID as of 2/23/2019. The coding regions were aligned and 223 mutations were identified with 68 synonymous and 115 nonsynonymous changes. The directionality of changes was inferred based on the RaTG13 sequence. Frequency spectra of both synonymous and nonsynonymous changes are skewed. While the former shows excess of both high and low frequency mutations, the latter mainly exhibits an excess of low frequency changes (Fig. 1a). The excess of low frequency mutations is consistent with the recent origin of SARS-CoV-2 [19]. Both population reduction and positive selection can increase high frequency mutations [20, 21]. However, the first scenario is contradicted by the recent origin of the virus. If positive selection has been operating, we would expect an excess of high frequency non-synonymous as well as synonymous changes. Furthermore, the ratio of nonsynonymous to synonymous changes is 2.46 (138/56) among singleton variants, but only 1.42 (17/12) among non-singletons. Both of these observations suggest that the majority of amino acid-altering mutations are selected against, with no positive selection in evidence.

**Figure 1.**
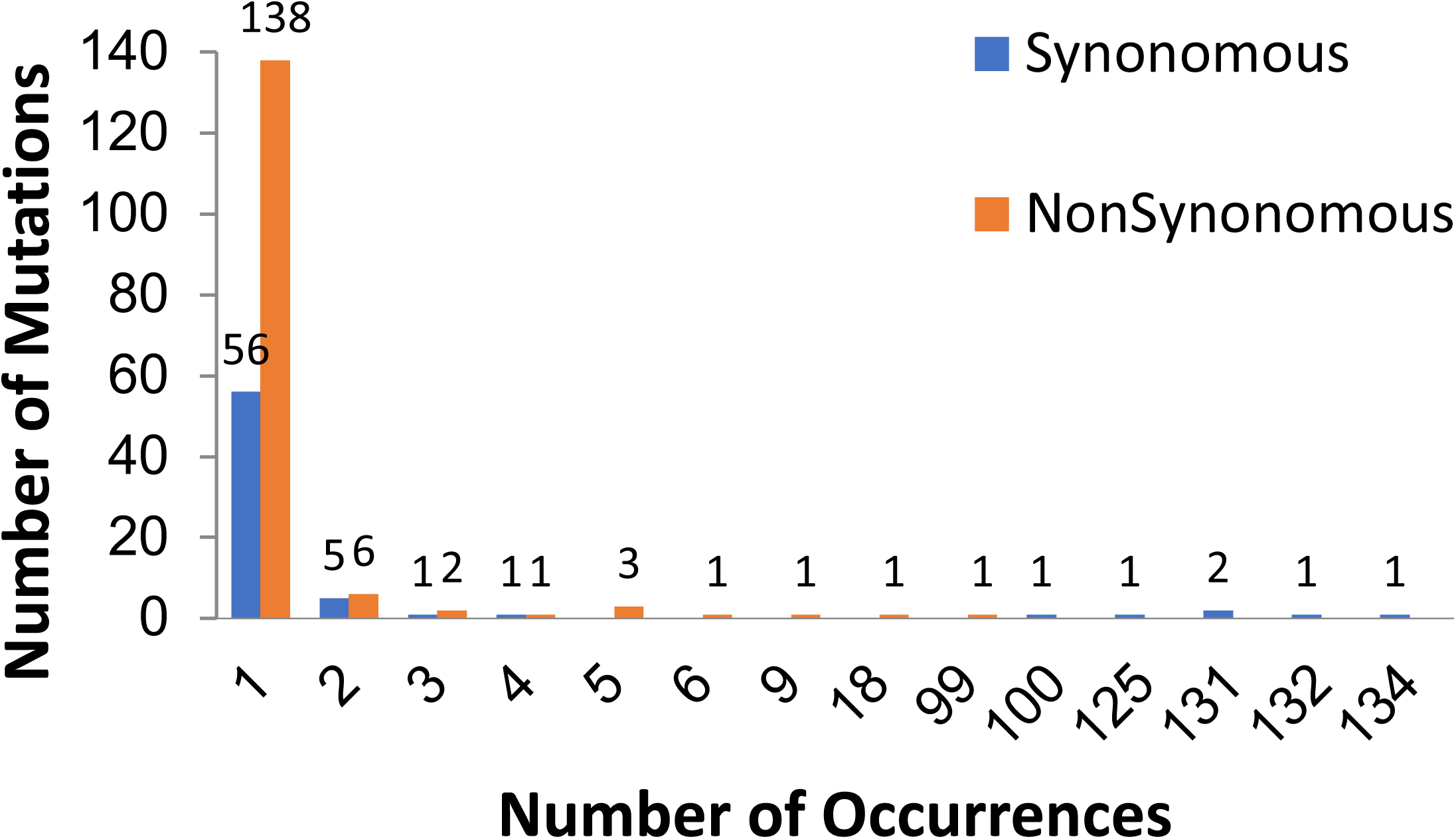
Frequency spectra of SARS-CoV-2. (A) The direction of changes was based on outgroup comparison with RaTG13. (B) The direction of changes was cross-referenced with the haplotype network showing in Fig. 2

The skew of synonymous variants toward high frequency deserves further discussion, as it relates to the underlying force driving the SARS-CoV-2 outbreak. The puzzle is probably rooted in how high and low frequency mutations are inferred. The results shown in the Fig. 1a are based on an outgroup comparison. The divergence at synonymous sites between SARS-CoV-2 and RaTG13 is 17%, approximately 3-fold greater than between humans and rhesus macaques [22]. With such high level of divergence, the possibility of multiple substitutions cannot be ignored, especially since substitution in the coronavirus genome is strongly biased toward transitions (see above). Indeed, among all non-singleton mutations listed in Table 2, 62% of the changes are C-T transitions.

**Table 2.** List of non-singleton mutations of SARS-CoV-2

|  | Genome position | Gene | RaTG13 | Pangolin_2<br>017 | Pangolin_2<br>019 | Major allele | Minor allele | amount of change |  |  |
| --- | --- | --- | --- | --- | --- | --- | --- | --- | --- | --- |
| Nonsynonymous |  |  |  |  |  |  |  | I | II |  |
| A | 614 | *orf1ab* | G | G | G | G | A | 2 |  | H116Q |
| B | 1190 | *orf1ab* | C | C | C | C | T | 3 |  | P308S |
| C | 5084 | *orf1ab* | A | A | A | A | G | 2 |  | A1606T |
| D | 9438 | *orf1ab* | C | C | C | C | T | 3 |  | T3058I |
| E | 11083 | *orf1ab* | G | T | G | G | T | 9 |  | L3606F |
| F | 18488 | *orf1ab* | T | T | T | T | C | 2 |  | I6074V |
| G | 21707 | *S* | C | C | N/A | C | T | 5 |  | H48Y |
| H | 22661 | *S* | G | G | G | G | T | 5 |  | V366F |
| I | 26144 | *orf3* | G | G | G | G | T | 18 |  | G251V |
| J | 27147 | *M* | G | G | G | G | C | 2 |  | I208T |
| K | 28077 | *orf8* | G | G | G | G | C | 4 |  | V61L |
| L | 28144 | *orf8* | C | C | C | T | C | 99 | 38 | L84S |
| M | 28854 | *N* | C | C | C | C | T | 5 |  | S194L |
| N | 28878 | *N* | G | G | G | G | A | 6 |  | S202N |
| O | 29019 | *N* | A | A | A | A | T | 2 |  | D249H |
| P | 29303 | *N* | C | C | C | C | T | 2 |  | K343I |
| Synonymous |  |  |  |  |  |  |  |  |  |  |
| $\alpha$ | 2662 | *orf1ab* | C | T | T | C | T | 3 | | C2397T |
| $\beta$ | 8782 | *orf1ab* | T | T | T | C | T | 100 | 37 | C8517T |
| $\gamma$ | 10138 | *orf1ab* | T | T | T | C | T | 134 | 3 | C9873T |
| $\delta$ | 15324 | *orf1ab* | C | C | C | C | T | 2 | | C15059T |
| $\epsilon$ | 17373 | *orf1ab* | T | C | T | C | T | 132 | 5 | C17108T |
| $\zeta$ | 18060 | *orf1ab* | T | T | A | C | T | 131 | 6 | C17795T |
| $\eta$ | 18603 | *orf1ab* | T | T | C | T | A | 2 | | T18338C |
| $\theta$ | 23569 | *S* | A | C | A | T | C | 2 | | T2007C |
| ι | 23605 | *S* | N/A | N/A | N/A | T | G | 2 |  | T2043G |
| κ | 24034 | *S* | T | C | C | C | T | 131 | 6 | C2472T |
| λ | 24325 | *S* | A | A | A | A | G | 2 |  | A2763G |
| μ | 26729 | *M* | T | T | T | T | C | 4 |  | T207C |
| ν | 29095 | *N* | T | T | T | C | T | 125 | 12 | C822T |

I Number of changes was inferred by outgroup comparison only
II Number of changes was cross-referenced with the haplotype network of SARS-CoV-2, only numbers which are different from method I are shown.
*E: envelope; M: matrix; N: nucleocapsid; S: spike*

To get around the potential problem caused by multiple substitutions, we cross-referenced the course of changes using the SARS-CoV-2 haplotype network (Fig. 2) and phylogeny (S2 Fig). The two analyses yield very different pictures. For example, the highest frequency derived mutation in Table 2 is a C-T synonymous change at 10138 (marked γ in Fig 2 and Table 2). All three sequences from Singapore share the T nucleotide also found in the RaTG13 outgroup. Using the outgroup comparison, the C found in the rest of the human SARS-CoV-2 sequences is a derived mutation. However, the T at this position is restricted to genomes collected from Singapore on 2/4 and 2/6/2020 and not found in earlier samples. It is thus more sensible to infer that this T is a back mutation derived from C rather than an ancestral nucleotide. Another synonymous change at position 24034 occurred twice (C24034T) on different genomic backgrounds (marked κ in Fig 2). Although the outgroup sequence at this position is T, it is more likely that the C at this position is the ancestral nucleotide. We observed a number of such back or repeated mutations. An A-T nonsynonymous change at 29019 (D249H in nucleocapsid protein, marked O in Fig. 2) also occurred twice.

**Fig. 2.**
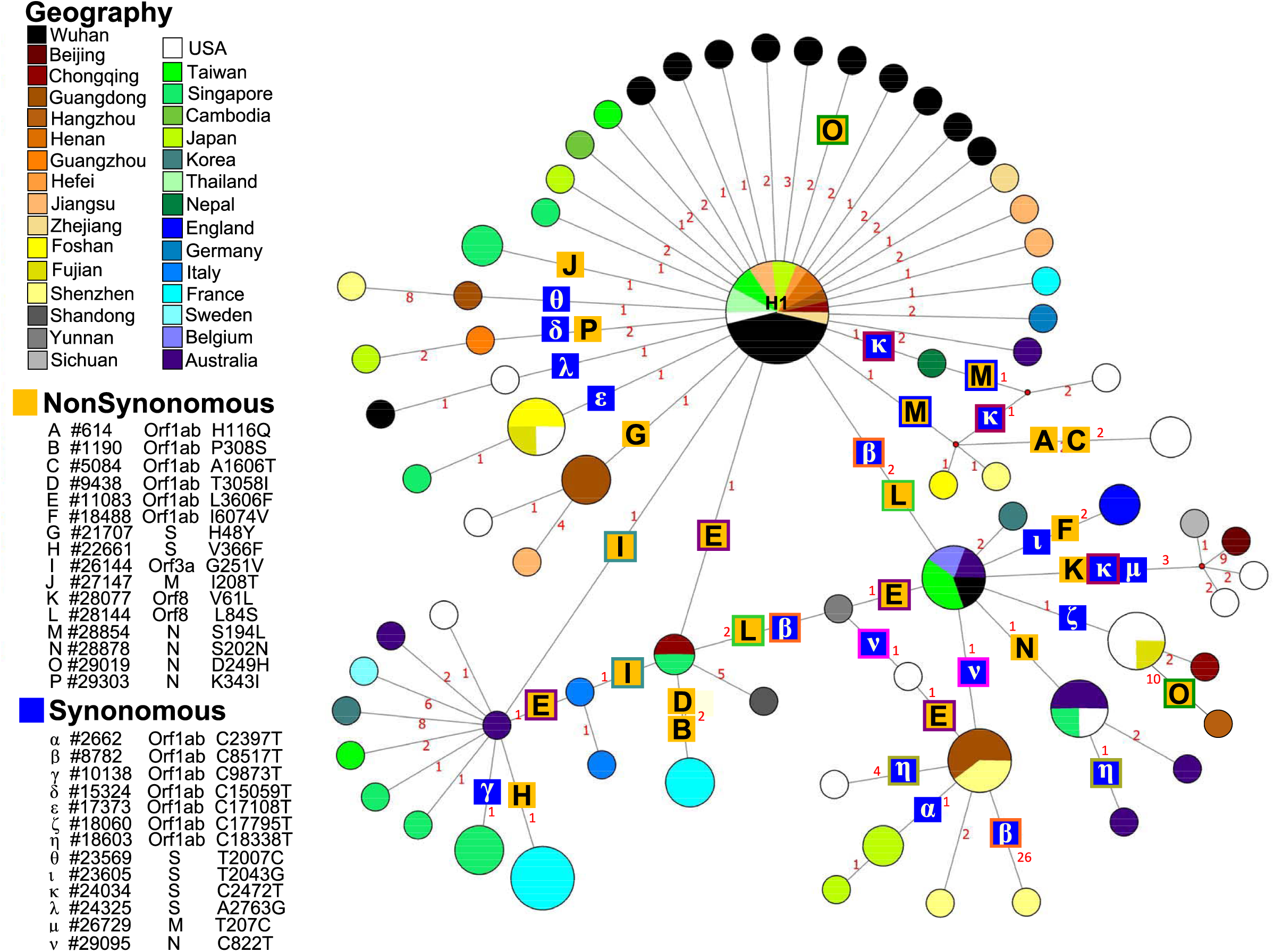
Haplotype network of SARS-CoV-2. Mutation types and numbers are given along the branch. Mutations that are involved in different evolutionary pathways or occurred more than once are enclosed. Also see Table 2 for comparison. Six genomes, EPI_ISL_ 408511, 408512, 410480, 408483, 407079, 407079, were excluded from this analysis because they contain too many ‘N’ in the sequences.

Repeated mutations may be caused by intergenomic recombination. Indeed, the result of four haplotype test suggested that at least two recombination events may have occurred between positions 8782 and 11083 and between 11083 and 28854. We noticed that a sequence isolated on 1/21/2020 from a patient in the United States (EPI_ISL_404253) exhibited Y (C or T) at both positions 8,782 and 28,144. Although, the possibility that two novel mutations might be occurred within this patient cannot be 100% ruled out, the alternative explanation that this patient may have been co-infected by two viral strains seems more plausible.

After cross-referencing with the haplotype network and the phylogeny, all mutations listed as high frequency in Table 2 and Fig. 1a were re-assigned to the other side of the frequency spectra. We only see an excess of singleton mutations, consistent with a recent origin of SARS-CoV-2 (Fig. 1b) and suggesting that the virus has mainly evolved under constraint.

Perhaps the most controversial case is the T-C change at position 28814 which alters Leucine (L) to Serine (S) in orf8 protein (L84S). Since both pangolin and RaTG13 have a C at this position (Table 2), Tang et al suggested that 84L is derived from 84S in the human virus [13]. The 84S was not discovered until 1/5/2020, by which time 23 SARS-CoV-2 genomes have been sampled. After the first appearance, its frequency gradually increased, reaching approximately 30% by 1/23/2020, suggesting that 84S may exhibit some advantage over 84L. If genomes carrying 84S were ancestral, it would be a challenge to explain its absence in early samplings. In addition, as mentioned above, C-T transitions are dominant in coronavirus evolution and multiple hits were observed in SARS-CoV-2 (Fig. 2). It is therefore possible that 28814C mutated to T after ancestral SARS-CoV-2 diverged from the common ancestor with RaTG13 and recently changed back to C. Finally, if 84L is indeed a derived haplotype and has rapidly increased in its frequency by positive selection, we would expect haplotypes carrying 84L to have accumulated more derived mutations than haplotypes with 84S. However, after correcting for mutational direction, the two haplotypes exhibited similar mutation frequency spectra (S3 Fig). The alternative hypothesis that 84S is a back mutation from 84L is more plausible.

### Selection pressure on SARS-CoV-2

In addition to L84S, a G-T transversion at 26114 which caused an amino acid change in orf3 protein (G251V) is also at intermediate frequency (Table 2). 251V was first seen on 1/22/2020 and gradually increased its frequency to 13% by our sampling date (Fig. 3). We note that the emergence of 84S in orf8 and 251V in orf3 are consistent with the lockdown of Wuhan on 1/23/2020. The former first appeared in early January, gradually increased its frequency, and reached a plateau around 1/23/2020. The latter showed up on 1/22/2020 and rapidly increased its frequency within two weeks.

**Fig. 3.**
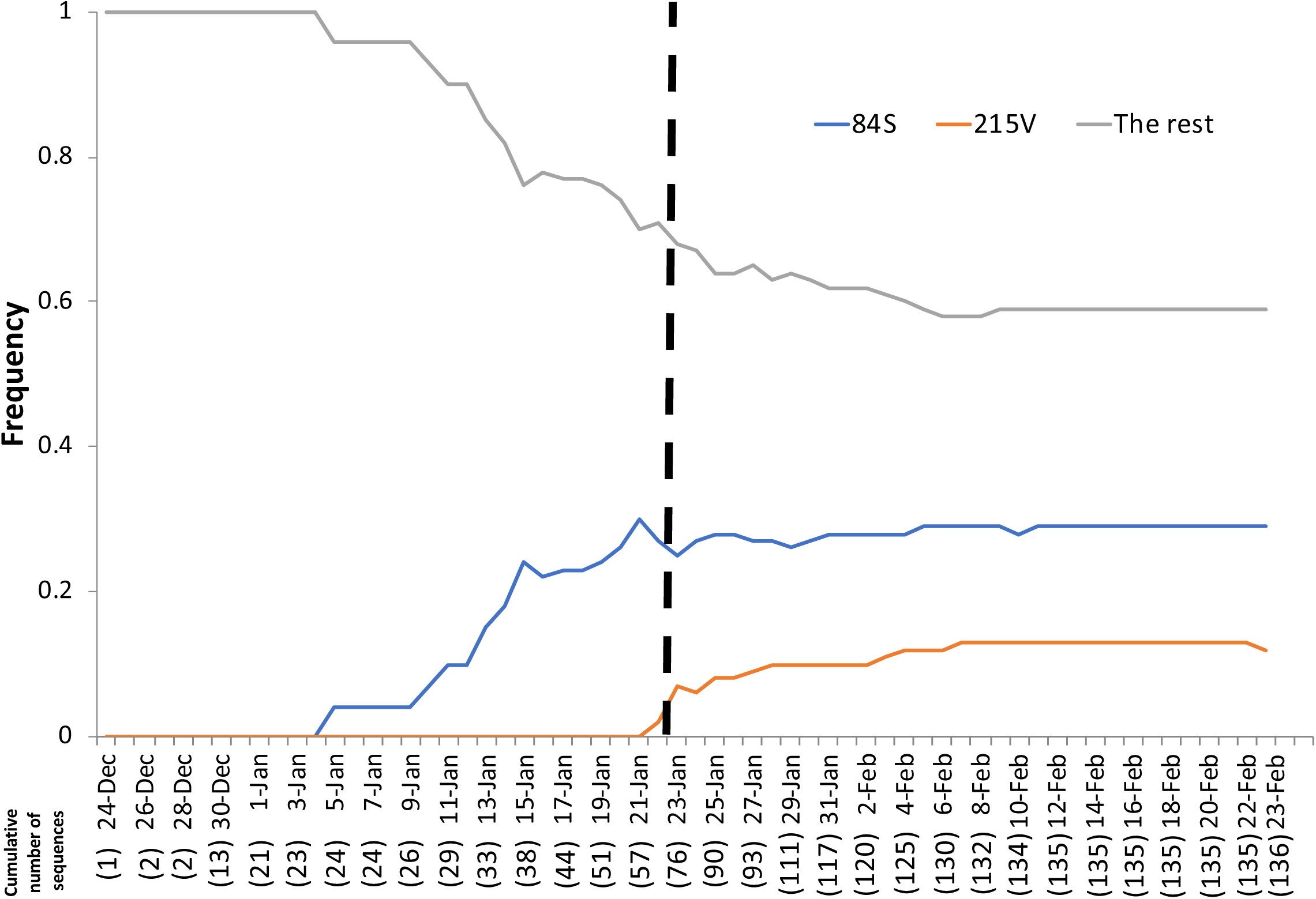
Mutation frequency of 84S is in orf8 and 215V is in orf3. The dashed line indicates the date of the Wuhan, China, lockdown.

Based on Fig. 3, we divided the sampling course into two epidemic episodes, from the first sampled sequence (12/24/2019) to before the lockdown of Wuhan (1/21/2020) and from 1/22/2020 to the date of the last sequence sampling (2/23/2020). The dN/dS of coding regions within the two episodes were estimated. As roughly 87% of mutations were singletons. Many of these are probably sequencing errors, affecting synonymous and nonsynonymous sites equally and inflating our dN/dS estimates. In addition, since dN/dS is already extremely small in SARS-CoV-2 (Table 1), such inflation would have a large effect on dN/dS estimates. We therefore excluded singletons from dN and dS estimation.

The dN/dS of *orf8* gene in episode I and II and *orf3* gene in episode II show strong signatures of positive selection, consistent with increase of 84S and 251V frequency during these periods, and may suggest a role of adaptation (Table 3). The overall dN/dS within each episode was 5-10 times higher than dN/dS between coronavirus genomes derived from different species (Table 1). The elevated dN/dS of SARS-CoV-2 is either due to its adaptation to human hosts or relaxation of selection. For a recently emerged virus, it is reasonable to expect operation of positive selection at the early stage. In that case, the dN/dS during episode I should be greater than during episode II [23, 24].

**Table 3.**
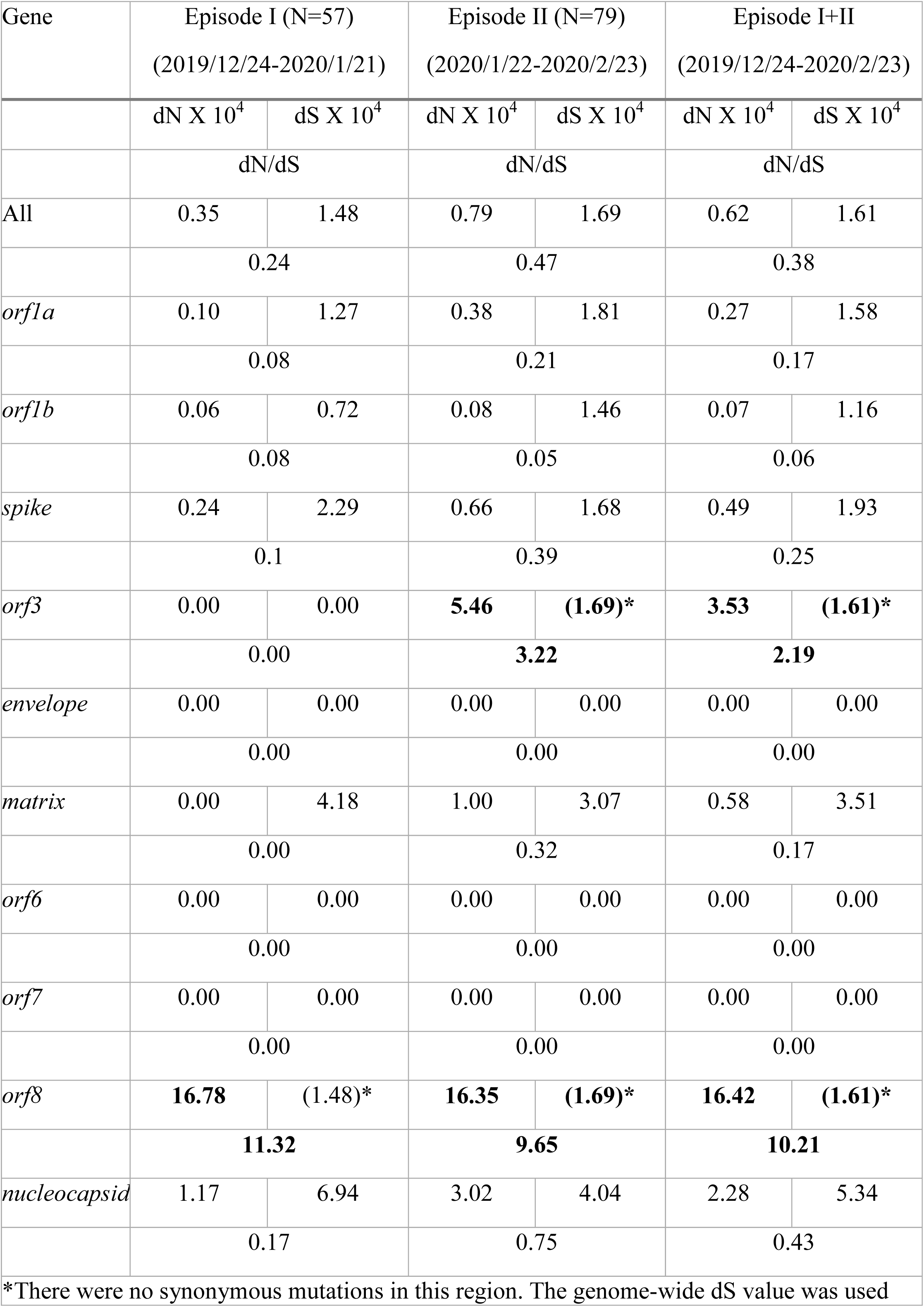
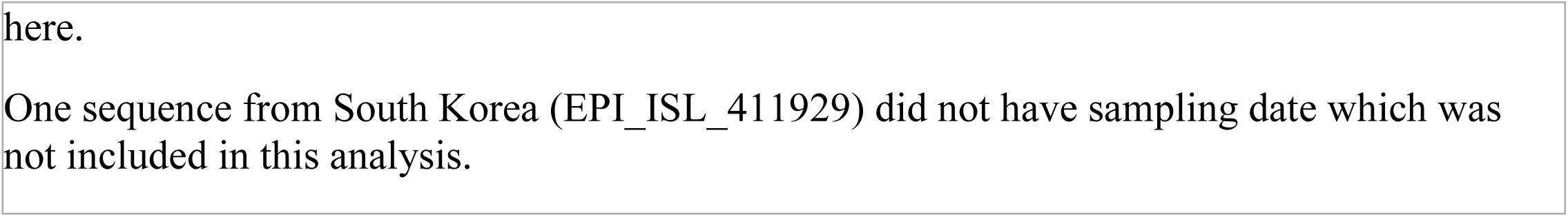
List of dN, dS, and dN/dS in coding regions of SARS-CoV-2 within two episodes

| Gene | Episode I (N=57)<br>(2019/12/24-2020/1/21) |  | Episode II (N=79)<br>(2020/1/22-2020/2/23) |  | Episode I+II<br>(2019/12/24-2020/2/23) |  |
| --- | --- | --- | --- | --- | --- | --- |
|  | dN X 10<sup>4</sup> | dS X 10<sup>4</sup> | dN X 10<sup>4</sup> | dS X 10<sup>4</sup> | dN X 10<sup>4</sup> | dS X 10<sup>4</sup> |
|  | dN/dS |  | dN/dS |  | dN/dS |  |
| All | 0.35 | 1.48 | 0.79 | 1.69 | 0.62 | 1.61 |
|  | 0.24 |  | 0.47 |  | 0.38 |  |
| *orf1a* | 0.10 | 1.27 | 0.38 | 1.81 | 0.27 | 1.58 |
|  | 0.08 |  | 0.21 |  | 0.17 |  |
| *orf1b* | 0.06 | 0.72 | 0.08 | 1.46 | 0.07 | 1.16 |
|  | 0.08 |  | 0.05 |  | 0.06 |  |
| *spike* | 0.24 | 2.29 | 0.66 | 1.68 | 0.49 | 1.93 |
|  | 0.1 |  | 0.39 |  | 0.25 |  |
| *orf3* | 0.00 | 0.00 | **5.46** | **(1.69)\*** | **3.53** | **(1.61)\*** |
|  | 0.00 |  | **3.22** |  | **2.19** |  |
| *envelope* | 0.00 | 0.00 | 0.00 | 0.00 | 0.00 | 0.00 |
|  | 0.00 |  | 0.00 |  | 0.00 |  |
| *matrix* | 0.00 | 4.18 | 1.00 | 3.07 | 0.58 | 3.51 |
|  | 0.00 |  | 0.32 |  | 0.17 |  |
| *orf6* | 0.00 | 0.00 | 0.00 | 0.00 | 0.00 | 0.00 |
|  | 0.00 |  | 0.00 |  | 0.00 |  |
| *orf7* | 0.00 | 0.00 | 0.00 | 0.00 | 0.00 | 0.00 |
|  | 0.00 |  | 0.00 |  | 0.00 |  |
| *orf8* | **16.78** | (1.48)* | **16.35** | **(1.69)\*** | **16.42** | **(1.61)\*** |
|  | **11.32** |  | **9.65** |  | **10.21** |  |
| *nucleocapsid* | 1.17 | 6.94 | 3.02 | 4.04 | 2.28 | 5.34 |
|  | 0.17 |  | 0.75 |  | 0.43 |  |

\*There were no synonymous mutations in this region. The genome-wide dS value was used
here.
One sequence from South Korea (EPI\_ISL\_411929) did not have sampling date which was not included in this analysis.

However, dN/dS was smaller in episode I than in episode II across the majority of the genome, suggesting that elevation of dN/dS is probably mostly due to the relaxation of selection. We further divided episode I into Ia and Ib, according to the appearance of 84S in orf8 protein on 1/6/2020. The genome-wide dN/dS values were 0.27 and 0.23 for episode 1a and 1b, respectively (S1 Table). Therefore, as shown in the frequency spectra, the signature of positive selection is weak at the early stage of the epidemic.

### The origin of SARS-CoV-2

The estimated mutation rate of SARS-CoV-2 is 2.4×10^−3^/site/year with 95% highest posterior density (HPD) of 1.5-3.3×10^−3^/site/year. The mutation rate at the third codon position is 2.9×10^−3^/site/year (95% HPD 1.8−4.0×10^−3^/site/year), which is in a good agreement with synonymous mutation rate of SARS-CoV, 1.67−4.67 × 10^−3^ /site/year [24]. SARS-CoV-2 is estimated to have originated on 12/11/2019 (95% HPD 11/13/2019−12/23/2019). We have to point out that the TMRCA estimation is strongly influenced by the genome sampling scheme. Since the earliest available genome was sampled on 12/24/2019 almost one month after the outbreak, the real origin of the current outbreak may actually be earlier than our estimation.

We estimated genetic variation, including the number of segregating sites, Watterson’s estimator of θ, and nucleotide diversity (π) of the SARS-CoV-2. Since both π and θ are estimators of 4Nu (N and u are the effective population size and mutation rate, respectively), they should be close to each other at the mutation-drift equilibrium [25]. Because θ is strongly influenced by rare mutations which are common during recent population expansion [14], it is a better estimator of genetic diversity for SARS-CoV-2. For example, when all samples are considered, θ (13.92 × 10^−4^) is approximately eight times higher than π (1.81 × 10^−4^, Table 4). Among samples collected from different locations, sequences from China exhibited higher genetic variation in terms of the number of segregating sites, θ and π, than the rest of the world combined, consistent with the observation that the outbreak originated in China, as the source populations are expected to exhibit higher genetic variation than derived populations [25].

**Table 4.** Nucleotide diversity of SARS-CoV-2 across geographic regions

| Sample origin | Sample size | S | $\theta \times 10^{-4}$ | $\pi \times 10^{-4}$ |
| --- | --- | --- | --- | --- |
| Total | 137 | 223 | 13.92 | 1.81 |
| China | 64 | 157 | 11.38 | 2.10 |
| Wuhan | 24 | 41 | 2.76 | 1.16 |
| Rest of China | 40 | 119 | 9.59 | 2.62 |
| Rest of the World | 73 | 81 | 5.71 | 1.52 |
| USA | 17 | 28 | 2.84 | 1.71 |
| Rest of the World excluding USA | 56 | 62 | 4.63 | 1.43 |

S: Number of segregating sites.
$\theta$ : Nucleotide diversity based on Watterson (29)
$\pi$ : Nucleotide diversity based on Nei and Li (30)

The haplotype network also supports this notion (Fig. 2). Usually, ancestral haplotypes have a greater probability of being in the interior, have more mutational connections, and are geographically more widely distributed. The H1 haplotype is at the center of the network and is found in four countries and many places in China. In addition, a large portion of haplotypes is directly connected to H1. Therefore, it is likely that H1 is the ancestral haplotype. As 45% of H1 are found in Wuhan, this location is the most plausible origin of the ongoing pandemic.

## DISCUSSION

A close relationship between SARS-CoV-2 and pangolin_2019 at the amino acid level in the RBD region of the spike protein might be due to recent recombination [15, 16], data contamination, or convergent evolution. Since recent recombination and DNA contamination should affect synonymous and nonsynonymous sites equally, they can be convincingly rejected as great divergence at synonymous sites was observed in spite of similar amino acid sequences between the two genomes. While genotypic convergence may be observed in viruses repeatedly evolving under particular conditions, such as drug resistance and immune escape [26-29], it is otherwise rare. For adaptations that do not involve highly specialized conditions, divergent molecular pathways may develop and genotypic convergence would not be expected [30]. For example, SARS-CoV and SARS-CoV-2 both use the spike protein to bind human ACE2 [2], but five out of six critical amino acids within the RBD are different between these two viruses [17]. Since the SARS-CoV-2 and pangolin_2019 have diverged at about 47% of synonymous sites and infect different hosts, the idea that they share five out of six critical amino acids within RBD through convergent evolution seems far-fetched.

We therefore hypothesize that, instead of convergent evolution, the similarity of RBD between SARS-CoV-2 and pangolin_2019 was caused by an ancient inter-genomic recombination. Assuming a synonymous substitution rate of 2.9×10^−3^/site/year, the recombination was estimated to have occurred approximately 40 years ago (95% HPD: 31-69 years; divergence time (t) = divergence (dS)/(substitution rate x 2 × 3), considering dS in RBD is 3-fold of genome average). The amino acids in the RBD region of the two genomes have been maintained by natural selection ever since, while synonymous substitutions have been accumulated. If this is true, SARS-CoV-2 may have circulated cryptically among humans for years before being recently noticed.

The ancient origin of SARS-CoV-2 is supported by its lack of a signature of adaptive evolution as shown by frequency spectra and dN/dS in samples from the recent epidemic. For a recently acquired virus, rapid evolution and a strong signature of positive selection are expected. For example, during its short epidemic in 2002-2003, several rounds of adaptive changes have been documented in SARS-CoV genomes [23, 24]. After adapting to its host, the virus may evolve under purifying or relaxed selection, exactly as we see in SARS-CoV-2. Therefore, it is important to sequence samples from the early outbreak and to examine hospital archives for the trace of SARS-CoV-2 ancestors. This information not only can help us to understand the evolutionary path of this virus but also unravel the critical steps for it to achieve effective spreading in humans.

In addition to the RBD, the SARS-CoV-2 spike protein also contains a small insertion of a polybasic cleavage site which was thought to be unique within the B lineage of betacoronaviruses [17]. However, a recent analysis of bats collected from Yunnan, China, identified a similar insertion in a sequence, RmYN02, closely related to SARS-CoV-2, providing strong evidence that such seemingly sorcerous site insertions can occur in nature [11]. Both the polybasic cleavage site in RmYN02 and RBD in pangolin_2019 suggest that, like with SARS-CoV [6], all genetic elements required to form SARS-CoV-2 may have existed in the environment. More importantly, they can be brought together by frequent intergenomic recombination (see Result). Nature never runs out of material to create new pathogens. It is not whether but when and where the next epidemic will occur.

There is a heated debate about the evolutionary forces influencing the trajectory of the L84S mutation in orf8 protein (http://virological.org/t/response-to-on-the-origin-and-continuing-evolution-of-sars-cov-2/418). While Tang et al. considered Serine is the ancestral amino acid [13], we present evidence that it is a back mutation. The majority of sequences in Wuhan were sampled before early January 2020 and most genomes carrying 84S were found outside Wuhan after middle to late January 2020. The discrepancy in time and space impedes the effort to resolve the debate. It would require more sequences from the early stage of the epidemic to settle this issue. Regardless of its ancestral or derived status, we hypothesize that 84S may confer some selective advantage. Unless the sampling scheme is deliberately skewed, it is difficult to explain such dramatic frequency gain of 84S, from 0 to ∼30% in two weeks. Oddly, its frequency ceased to increase after 1/23/2020, when Wuhan was locked down. This coincidence prompts us to consider the effect of social distancing on virus transmission. Another line of evidence comes from the frequency increase of 215V in orf3 protein. The 215V first appeared on 1/22/2020 and rapidly increased its frequency within two weeks.

Several studies suggested that the orf8 protein may function in viral replication, modulating endoplasmic reticulum stress, inducing apoptosis, and inhibiting interferon responses in host cells (41-45 [31, 32-35]. During the SARS spread, frequency of several orf8 mutations fluctuated in accordance with different phases of the outbreak, suggesting that orf8 underwent adaptation during the SARS epidemic [24]. It is suggested that 84S may induce structural disorder in the C-terminus of the protein and may generate a novel phosphorylation target for Serine/Threonine kinases of the mammalian hosts [36].

SARS-CoV orf3 protein has been shown to activate NF-κB and the NLRP3 inflammasome and causes necrotic cell death, lysosomal damage, and caspase-1 activation. In addition, orf3 is required for maximal SARS-CoV replication and virulence. All of the above likely contributes to the clinical manifestations of SARS-CoV infection [37-39]. Therefore, these two mutations may have some functional consequences and be worth investigating further. By the time we prepared this manuscript, the 215V frequency ceased to increase. However, a parallel mutation has occurred in a different genomic background, further supporting the idea that this mutation may require further study.

## MATERIALS AND METHODS

### Data collection

137 complete SARS-CoV-2 genomes were downloaded from the Global Initiative on Sharing Avian Influenza Data (GISAID, https://www.gisaid.org/). Related coronavirus sequences, including those from five related bat sequences (RaTG13, HUK3-1, ZC45, ZXC-21, and GX2013), two pangolins (each from Guangdong (pangolin_2019) and Guangxi (pangolin_2017)), were downloaded from GenBank (https://www.ncbi.nlm.nih.gov/nucleotide/). Nucleotide positions and coding sequences (CDSs) of SARS-CoV-2 were anchored to the reference genome NC_045512. CDS annotations of other coronaviruses were downloaded from GenBank.

### Sequence analyses and phylogeny construction

CDSs were aligned based on translated amino acid sequences using MUSCLE v3.8.31 [40], and back-translated to their corresponding DNA sequences using TRANALIGN software from the EMBOSS package (http://emboss.open-bio.org/) [41]. Nucleotide diversity, including number of segregating sites, Watterson’s estimator of θ [42], and nucleotide diversity (π)[43], was estimated using MEGA-X [44]. MEGA-X was also used for phylogenetic construction. Phylogenetic relationships were constructed using the neighbor-joining method based on Kimura’s two-parameter model. Number of nonsynonymous changes per nonsynonymous site (dN) and synonymous changes per synonymous site (dS) among genomes were estimated based Li-Wu-Luo’s method [45] implemented in MEGA-X and PAML 4 [46].The RDP file for the haplotype network analyses was generated using DnaSP 6.0 [47] and input into Network 10 (https://www.fluxus-engineering.com/) to construct the haplotype network using the median joining algorithm. Four haplotype test implemented in DnaSp was applied to test for possible recombination event.

The mutation rate of SARS-CoV-2 and the time to the most recent common ancestor (TMRCA) of virus isolates were estimated by an established Bayesian MCMC approach implemented in BEAST version 1.10.4 [48]. The sampling dates were incorporated into TMRCA estimation. The analysis was performed using the HKY model of nucleotide substitution assuming an uncorrelated lognormal molecular clock [49]. We linked substitution rates for the first and second codon positions and allowed independent rates in the third codon position. We performed two independent runs with 3×10^8^ MCMC steps and the results were combined. Log files were checked using Tracer (http://beast.bgio.ed.ac.uk/Tracer). Effective sample sizes were >300 for all parameters.

## Acknowledgments

The authors thank those who contributed to sequence generation and sharing (The detail is listed in S2 Table). We also thank Chung-I Wu and Wen-Ya Ko for their constructive comments and suggestions. This work was supported by Ministry of Science and Technology, National Taiwan University, and National Taiwan University, College of Medicine, Taipei, Taiwan to HYW (105-2628-B-002-015-MY3, 107-2321-B-002-004-,NTU-109L7806, NSC-131-5), and partially by a grant form Biodiversity Research Center, Academia Sinica to SMC.

## Appendix Table

**Appendix Table 1.**
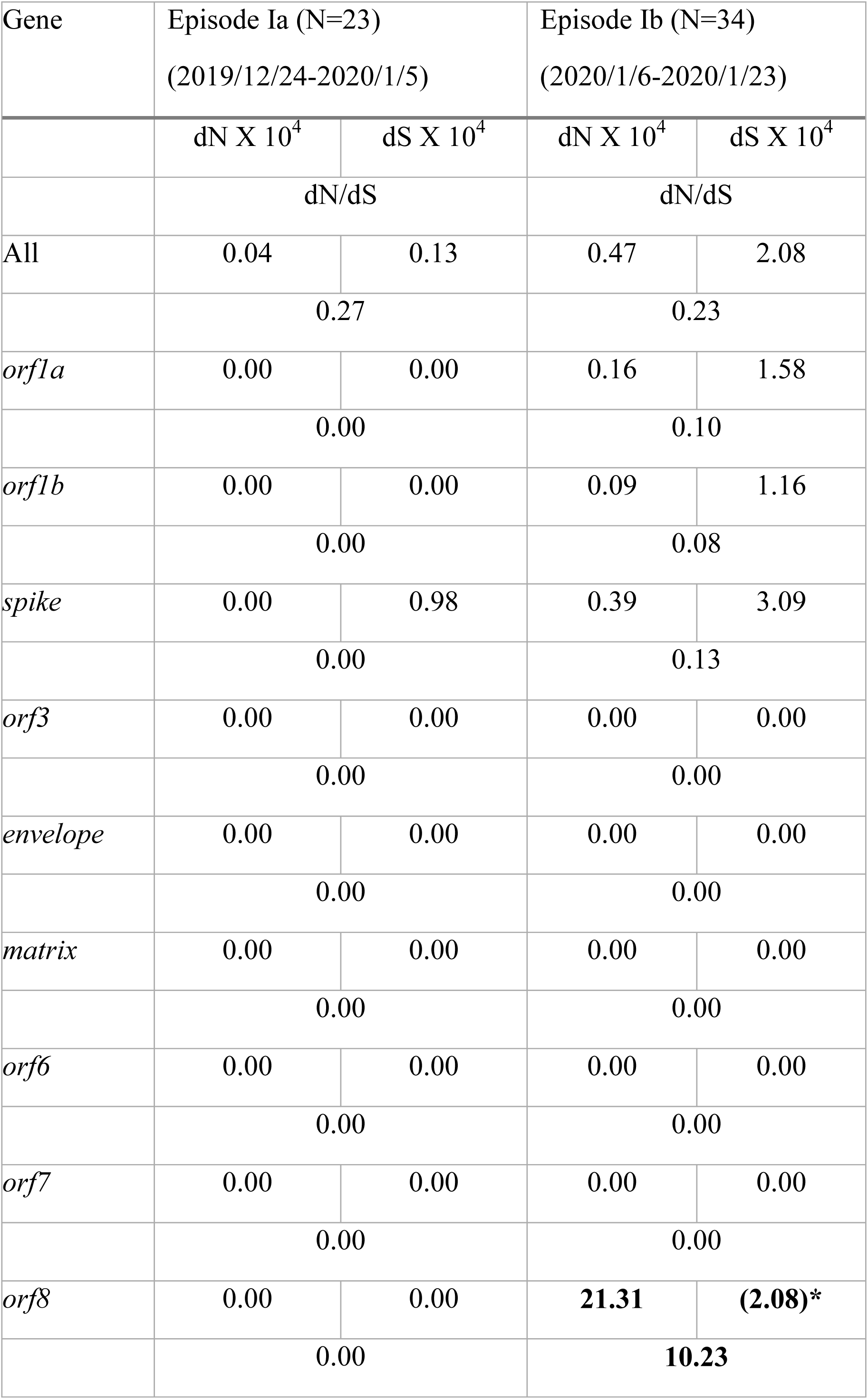
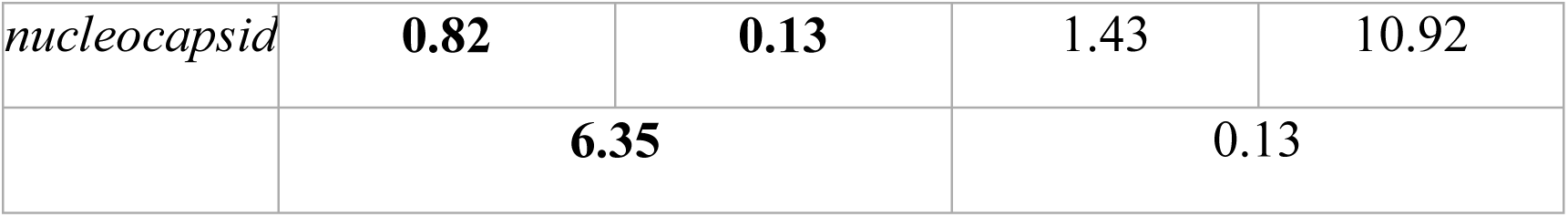
List of dN, dS, and dN/dS in coding regions of SARS-CoV-2 within episode Ia and Ib

| Gene | Episode Ia (N=23)<br>(2019/12/24-2020/1/5) |  | Episode Ib (N=34)<br>(2020/1/6-2020/1/23) |  |
| --- | --- | --- | --- | --- |
|  | dN X 10<sup>4</sup> | dS X 10<sup>4</sup> | dN X 10<sup>4</sup> | dS X 10<sup>4</sup> |
|  | dN/dS |  | dN/dS |  |
| All | 0.04 | 0.13 | 0.47 | 2.08 |
|  | 0.27 |  | 0.23 |  |
| *orf1a* | 0.00 | 0.00 | 0.16 | 1.58 |
|  | 0.00 |  | 0.10 |  |
| *orf1b* | 0.00 | 0.00 | 0.09 | 1.16 |
|  | 0.00 |  | 0.08 |  |
| *spike* | 0.00 | 0.98 | 0.39 | 3.09 |
|  | 0.00 |  | 0.13 |  |
| *orf3* | 0.00 | 0.00 | 0.00 | 0.00 |
|  | 0.00 |  | 0.00 |  |
| *envelope* | 0.00 | 0.00 | 0.00 | 0.00 |
|  | 0.00 |  | 0.00 |  |
| *matrix* | 0.00 | 0.00 | 0.00 | 0.00 |
|  | 0.00 |  | 0.00 |  |
| *orf6* | 0.00 | 0.00 | 0.00 | 0.00 |
|  | 0.00 |  | 0.00 |  |
| *orf7* | 0.00 | 0.00 | 0.00 | 0.00 |
|  | 0.00 |  | 0.00 |  |
| *orf8* | 0.00 | 0.00 | **21.31** | **(2.08)\*** |
|  | 0.00 |  | **10.23** |  |
| *nucleocapsid* | **0.82** | **0.13** | 1.43 | 10.92 |
|  | **6.35** |  | 0.13 |  |

## Appendix Figures

**Appendix Figure 1.**
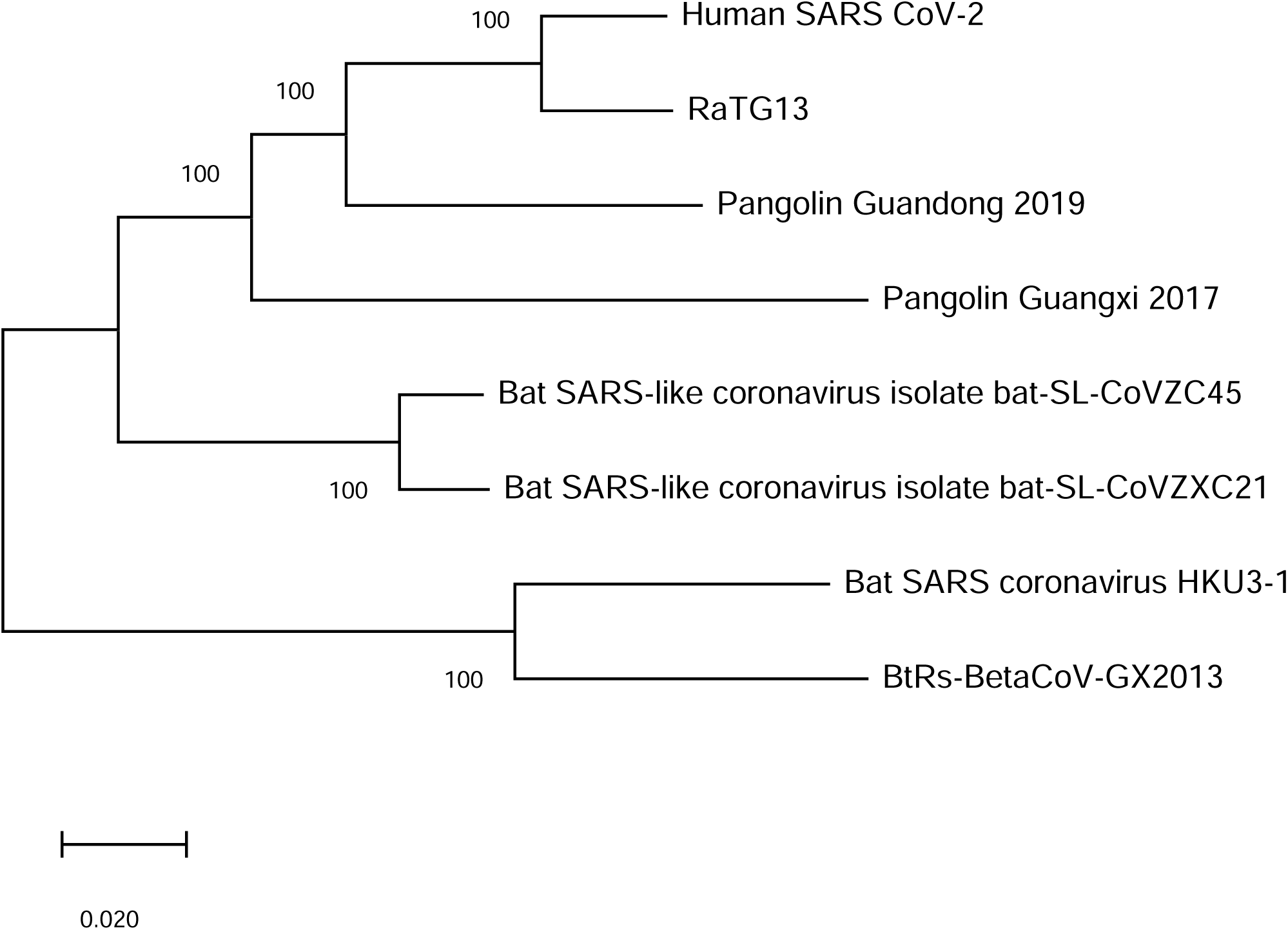
The neighbor-joining tree of SARS-CoV-2 related coronaviruses constructed by concatenating coding sequences based on the Kimura 2-parameter model implemented in MEGA-X.

**Appendix Figure 2.**
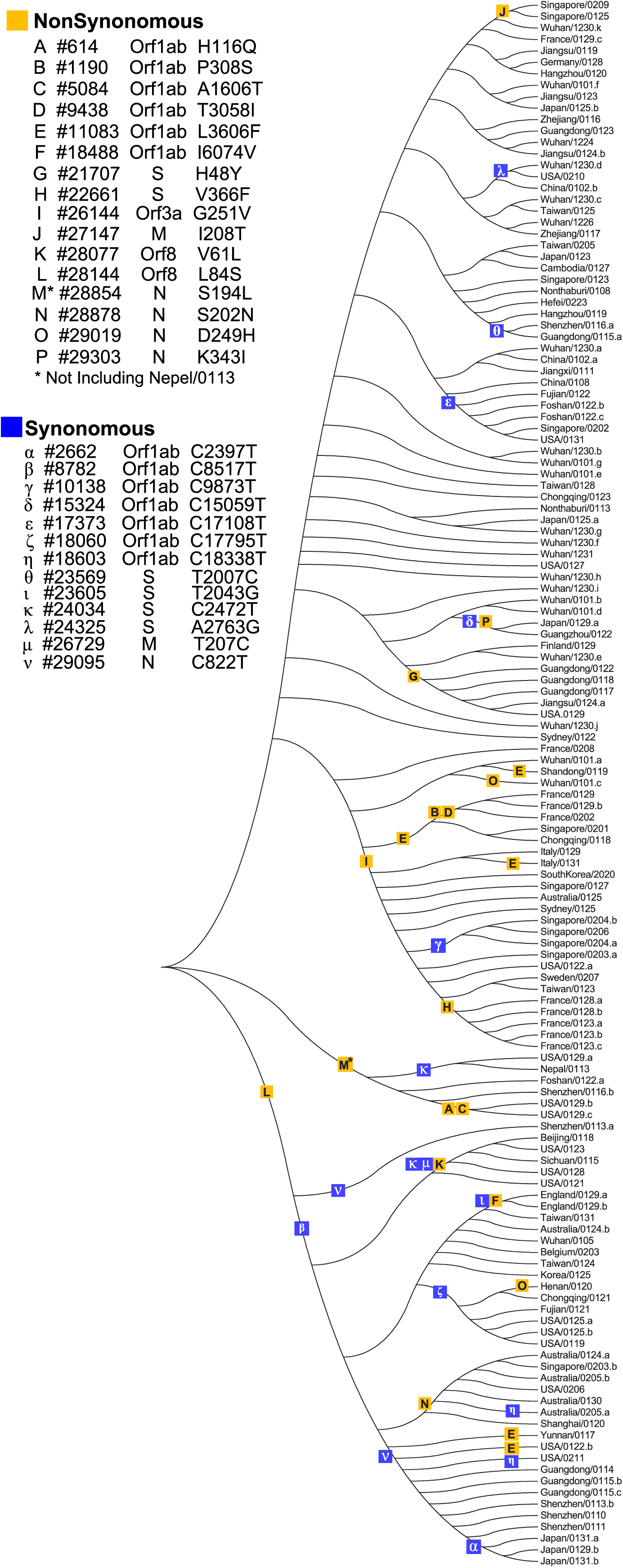
Unrooted neighbor-joining tree of SARS-CoV-2 constructed by concatenating coding sequences based on the Kimura 2-parameter model implemented in MEGA-X. Non-singleton changes are shown along the branches. The location of each sequence is given (above the slash) followed by its sampling date (below the slash). For multiple sequences sampled on the same date from the same location, the index, a, b, c, d, and etc. is given. Details are listed in Supplemental File 2.

**Appendix Figure 3.**
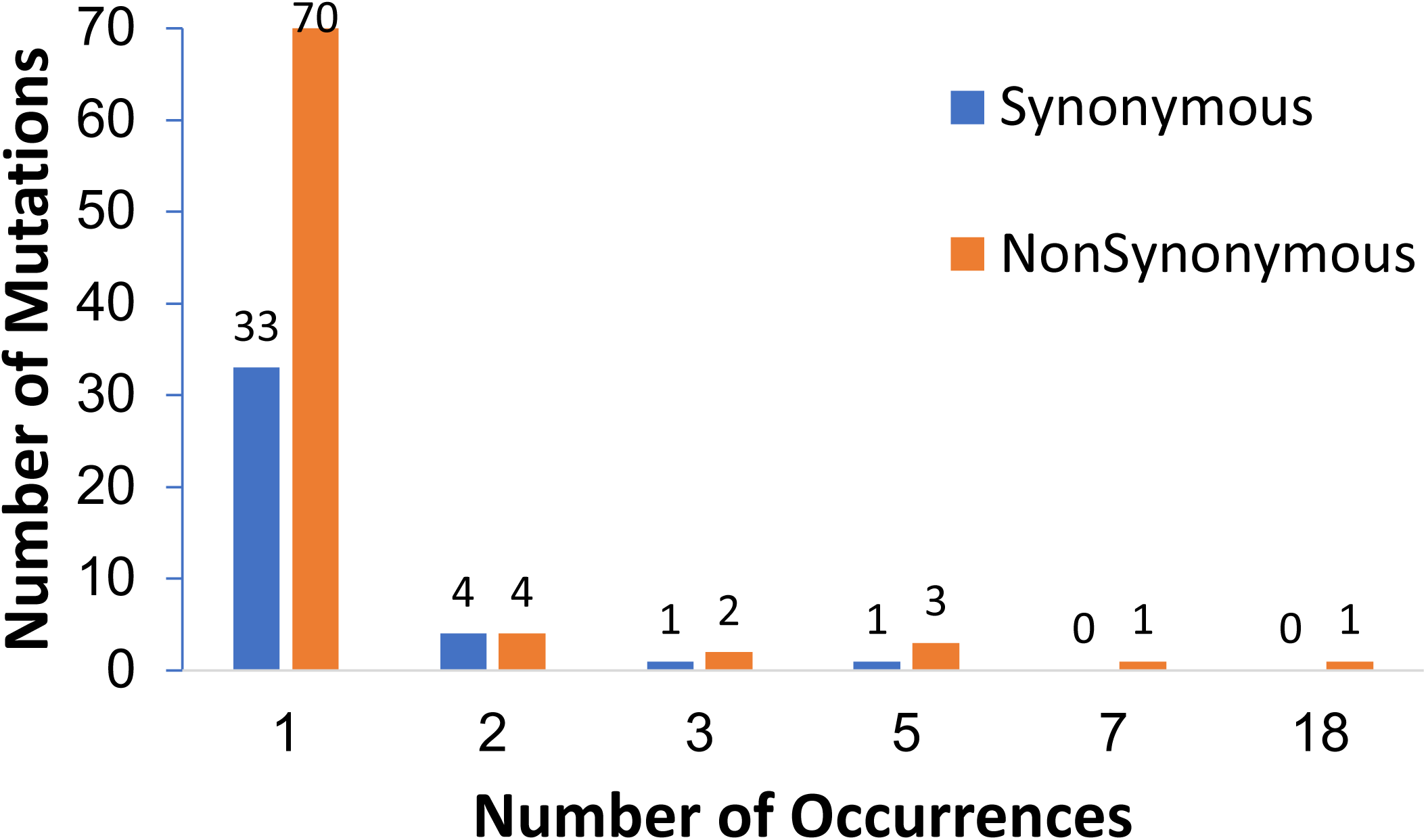
Frequency spectra of SARS-CoV-2 carrying 84L (n=98) (A) and 84S (n=39) (B) in orf8. The direction of changes was cross-referenced with the haplotype network shown in Fig. 2

**Appendix Figure 3.**
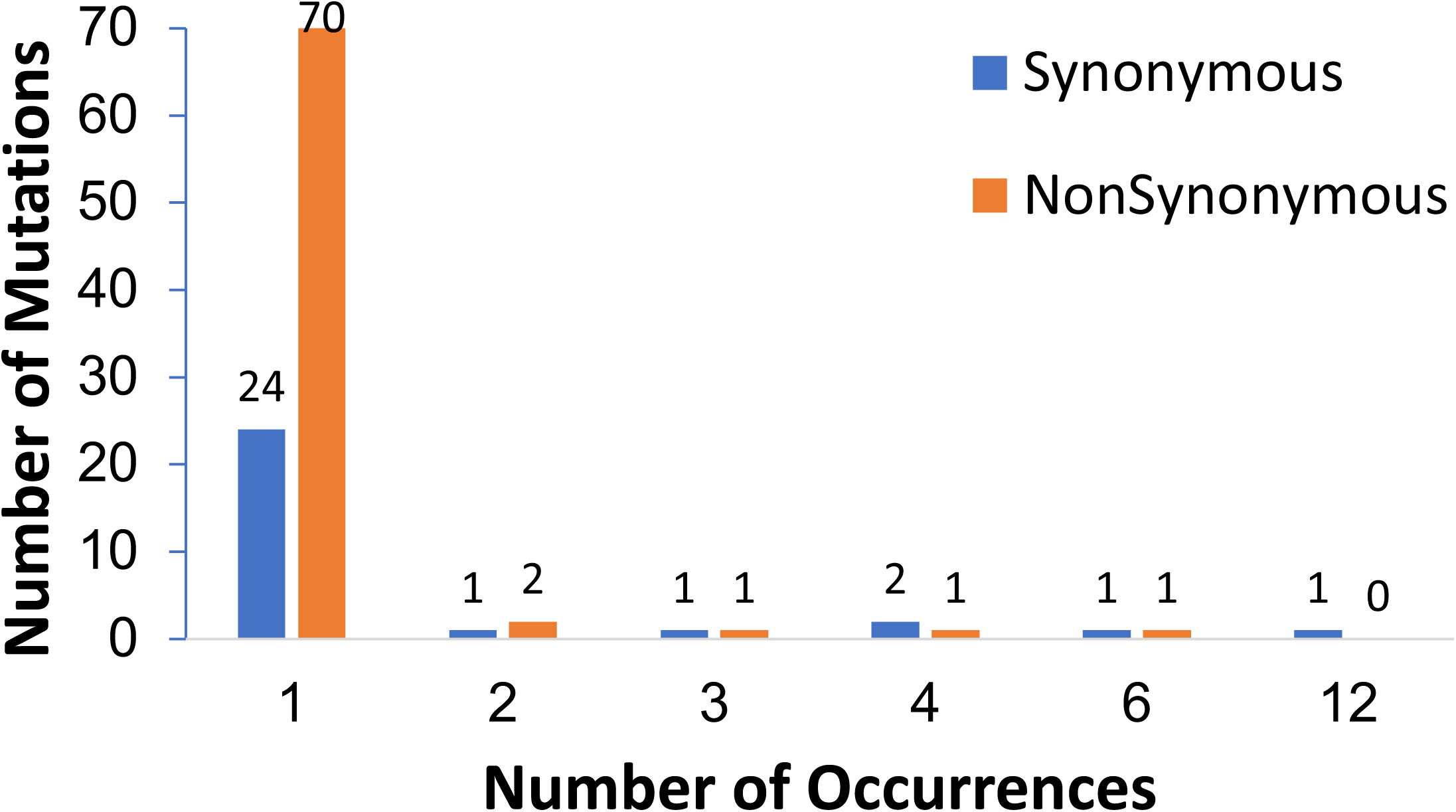
Frequency spectra of SARS-CoV-2 carrying 84L (n=98) (A) and 84S (n=39) (B) in orf8. The direction of changes was cross-referenced with the haplotype network shown in Fig. 2

---

**Figure 1.**
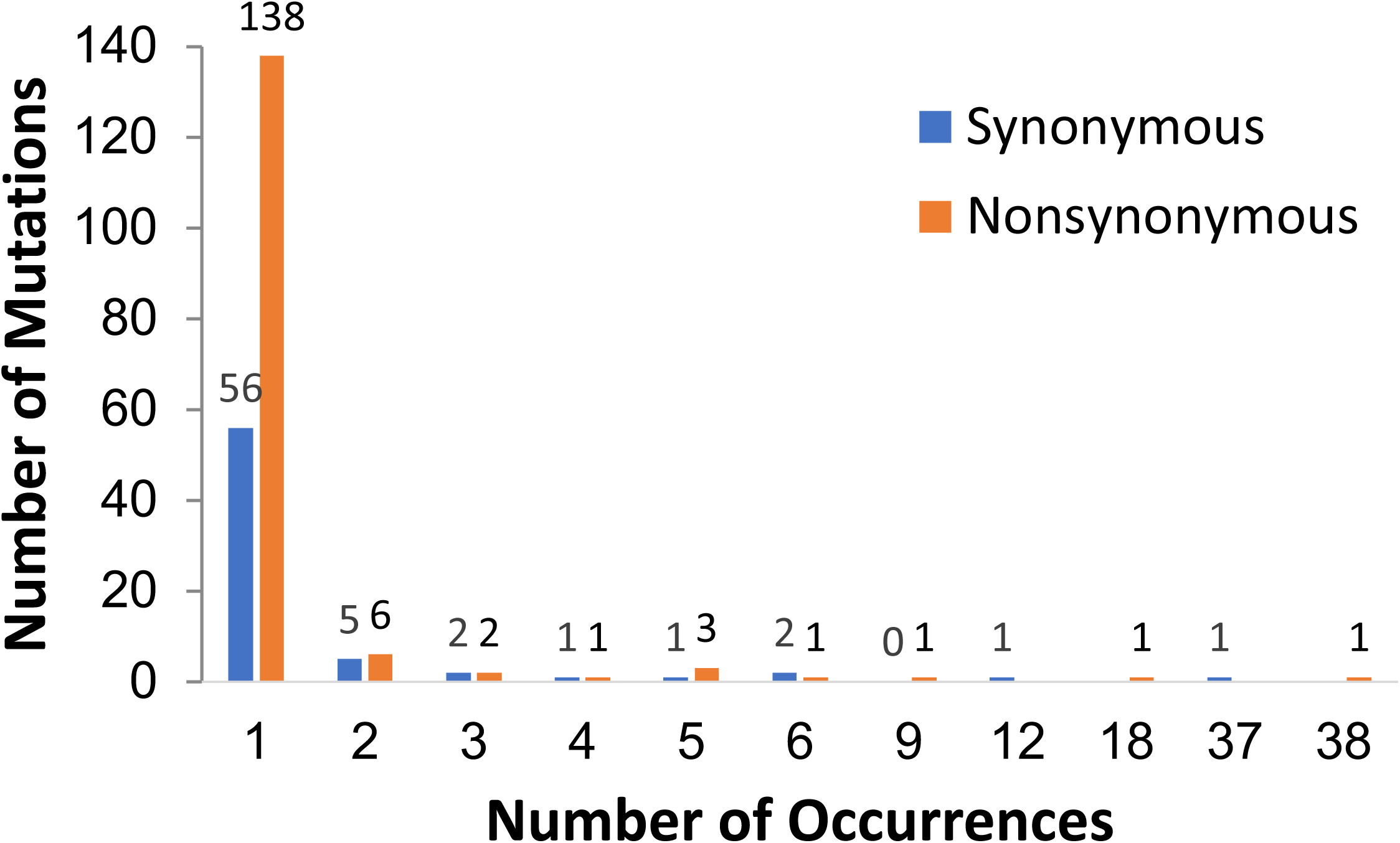
Frequency spectra of SARS-CoV-2. (A) The direction of changes was based on outgroup comparison with RaTG13. (B) The direction of changes was cross-referenced with the haplotype network showing in Fig. 2

